## Supplementary material for "p53-induced RNA-binding protein ZMAT3 inhibits transcription of a hexokinase to suppress mitochondrial respiration": Figure Supplement 1-5

**Figure 1- figure supplement 1**

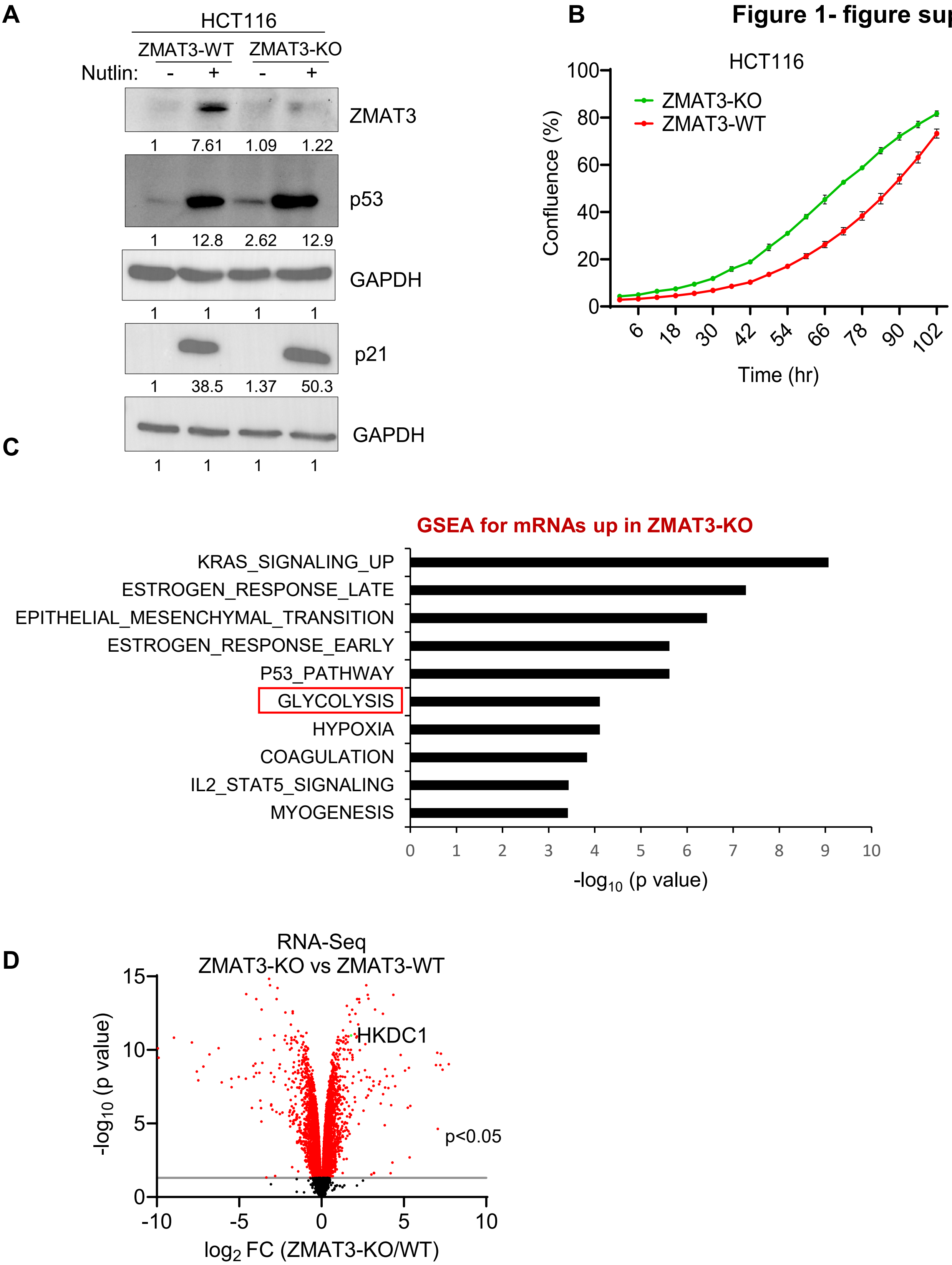

### Figure 2 – figure supplement 1

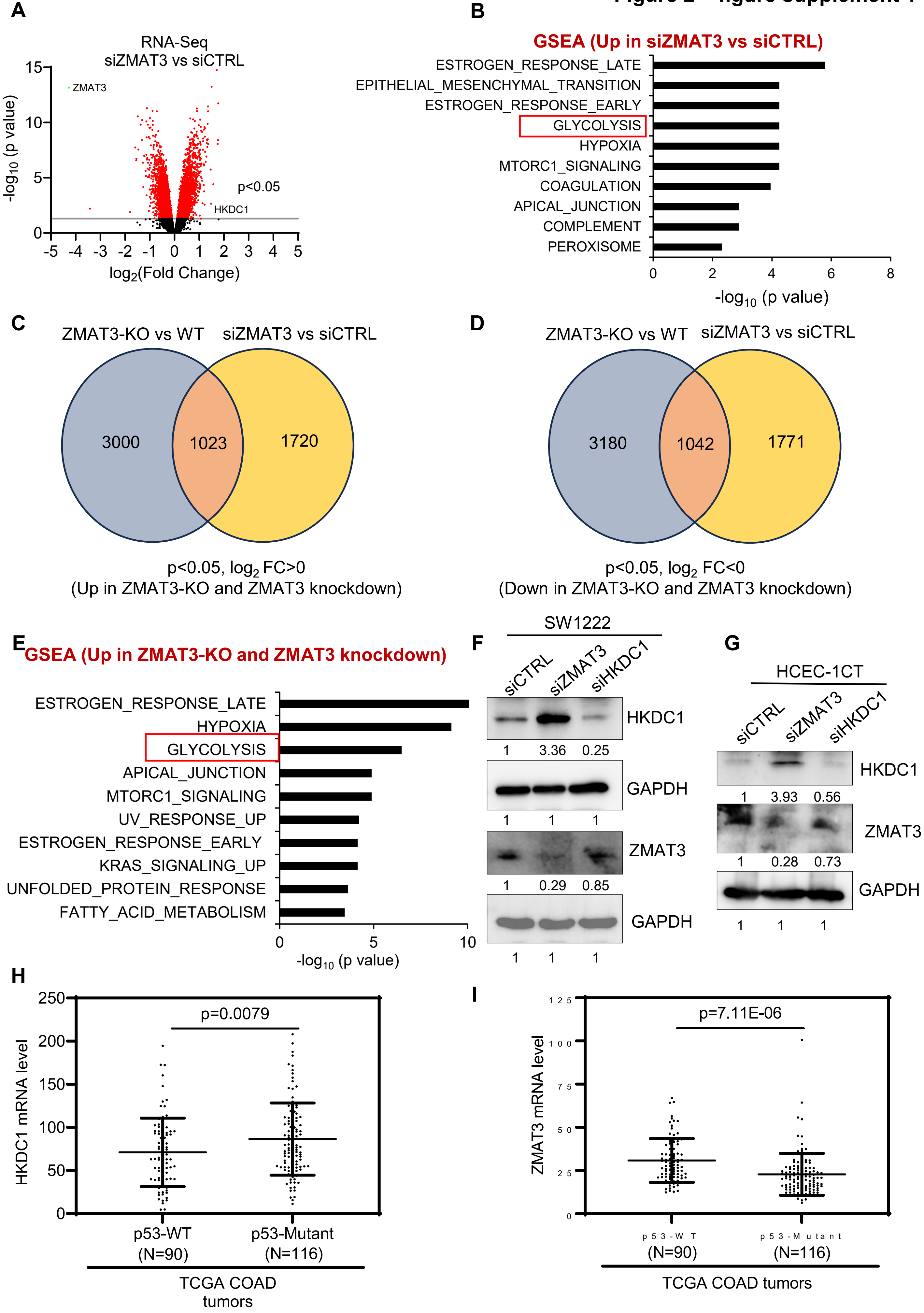

**Figure 3 – figure supplement 1**

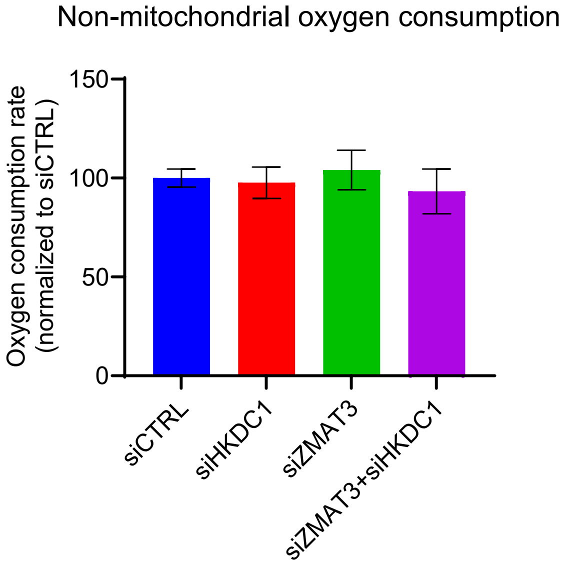

**Figure 4 – figure supplement 1**

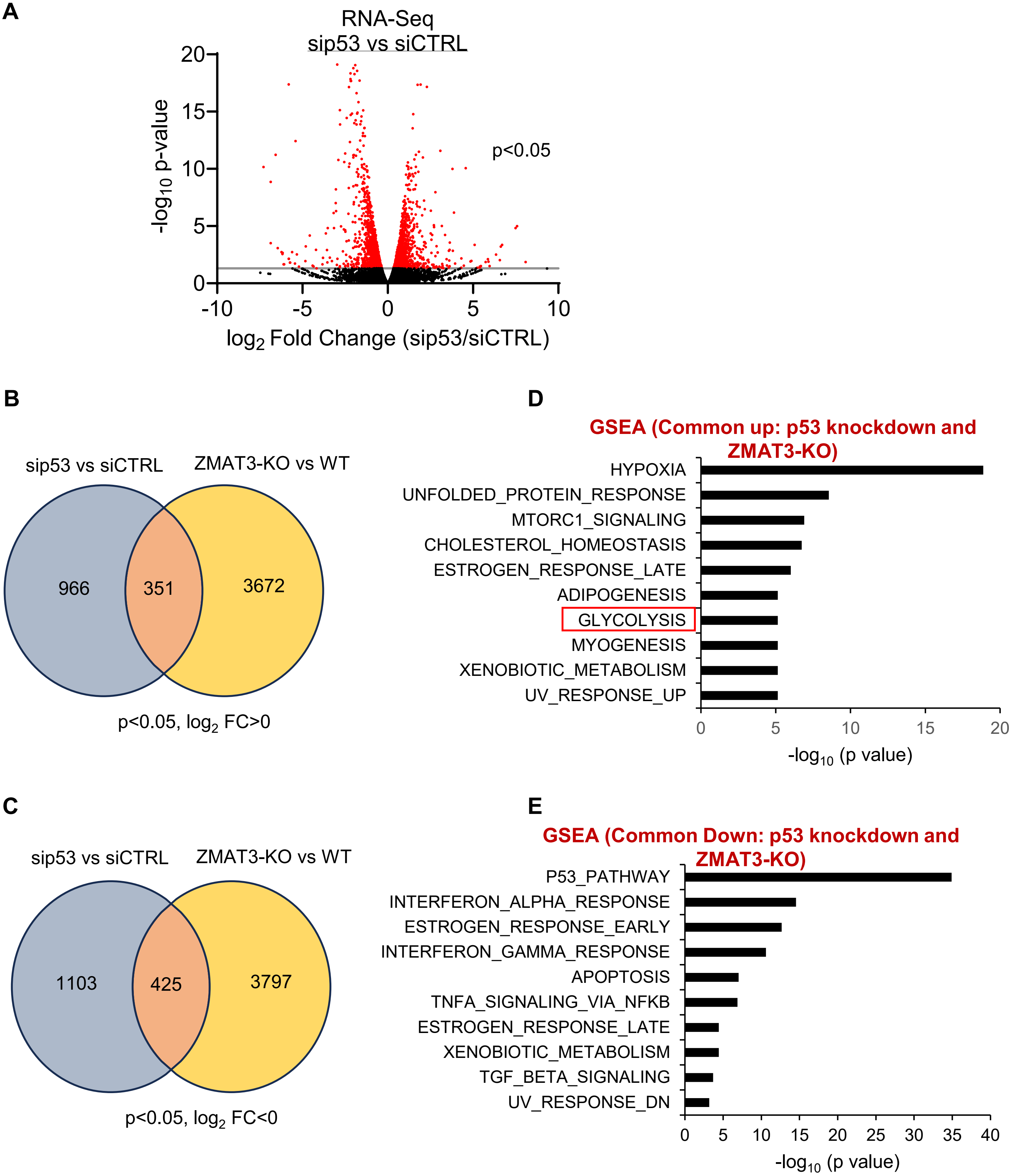

**Figure 5 – figure supplement 1**

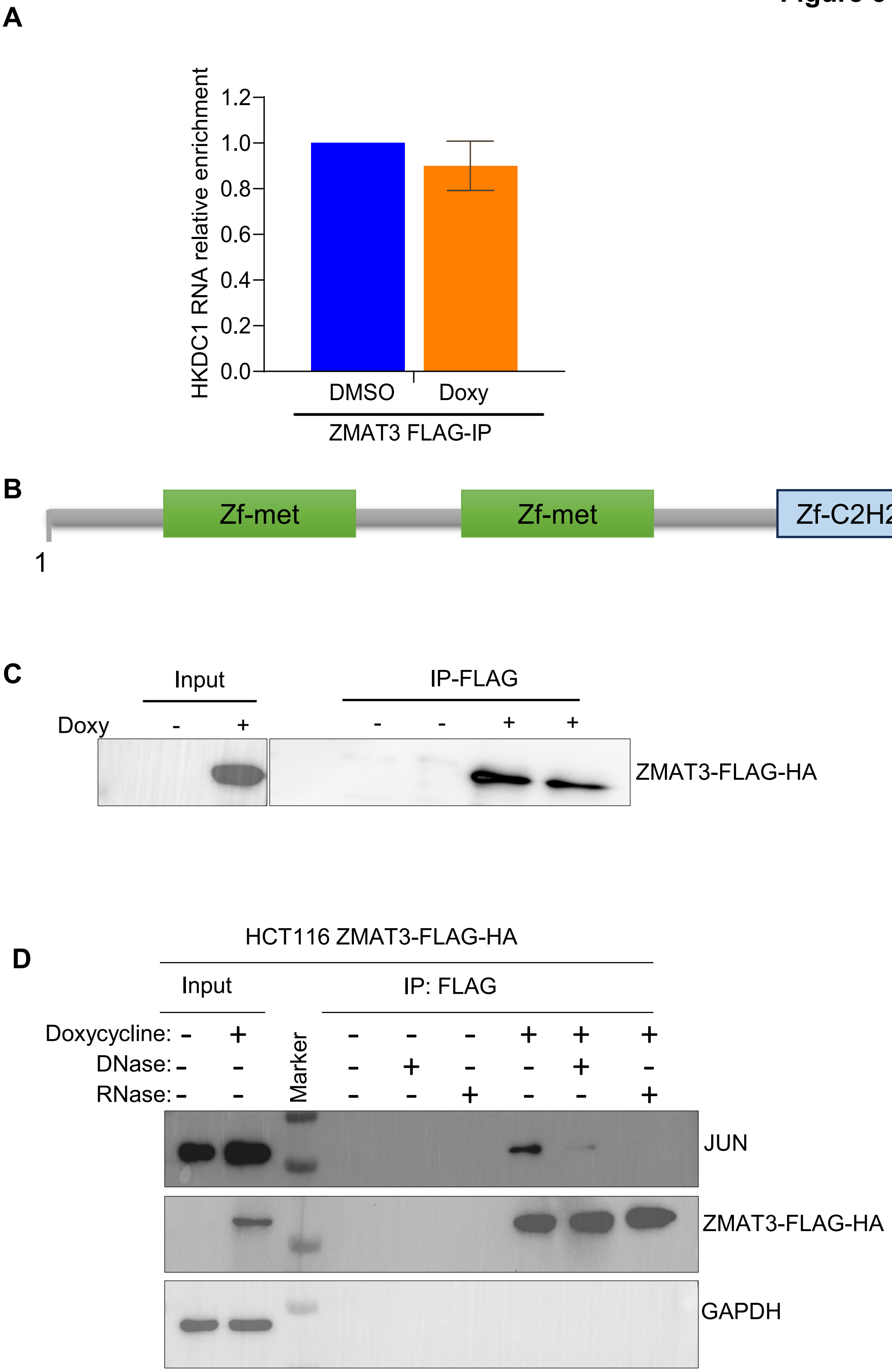

#### Figure 5 – figure supplement 2

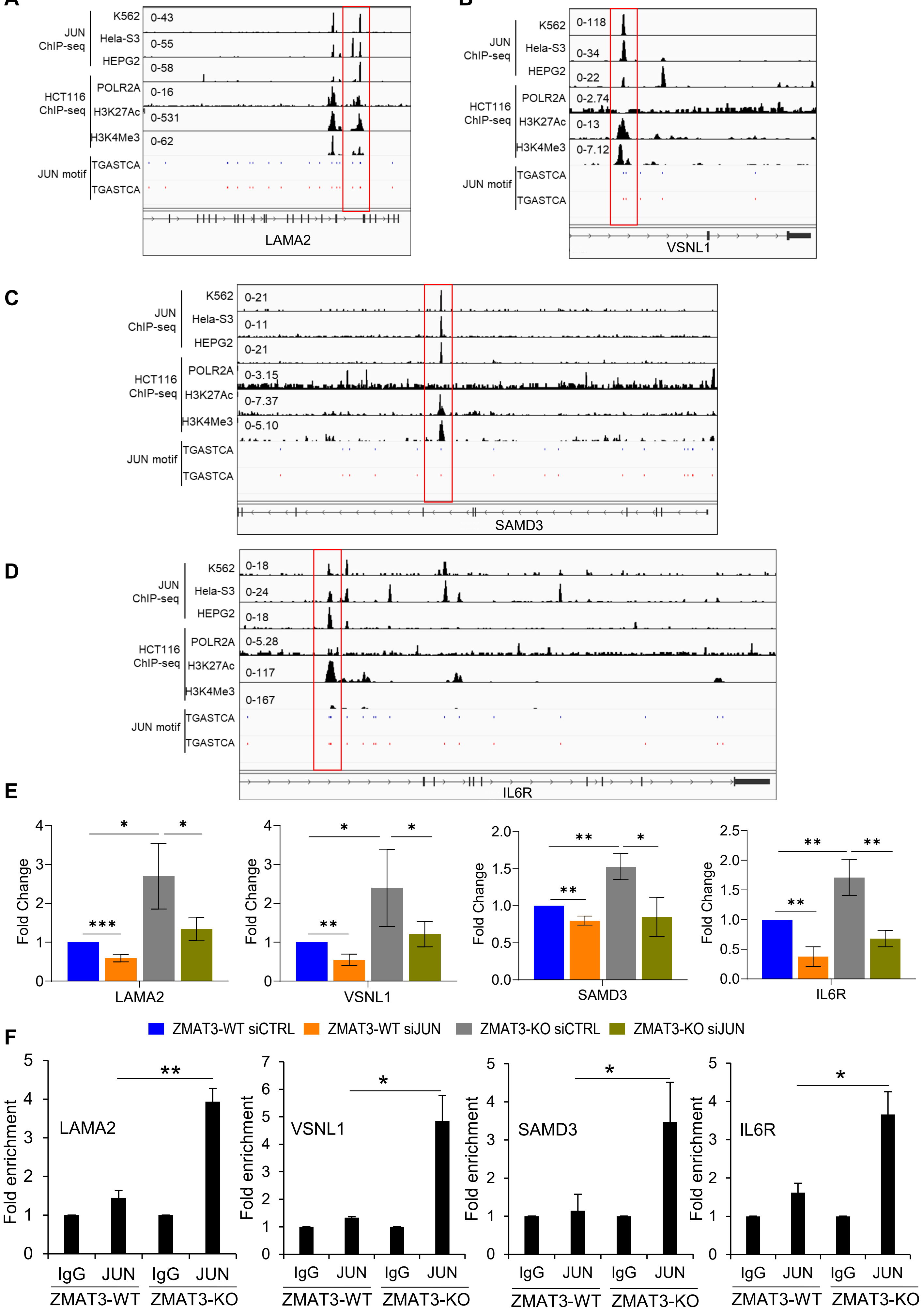
